# GuideMaker: Software to design CRISPR-Cas guide RNA pools in non-model genomes

**DOI:** 10.1101/2021.06.28.450164

**Authors:** Ravin Poudel, Lidimarie Trujillo Rodriguez, Christopher R. Reisch, Adam R. Rivers

**Affiliations:** Genomics and Bioinformatics Research Unit, USDA Agricultural Research Service, Gainesville, FL, 32608, USA; Department of Microbiology and Cell Science, Institute of Food and Agricultural Sciences, University of Florida, Gainesville, FL 326011, USA

**Keywords:** PAM, CRISPR-Cas, gRNA, HNSW

## Abstract

**Background:** CRISPR-Cas systems have expanded the possibilities for gene editing in bacteria and eukaryotes. There are many excellent tools for designing the CRISPR-Cas guide RNAs for model organisms with standard Cas enzymes. GuideMaker is intended as a fast and easy-to-use design tool for atypical projects with 1) non-standard Cas enzymes, 2) non-model organisms, or 3) projects that need to design a panel of guide RNAs (gRNA) for genome-wide screens.

**Findings:** GuideMaker can rapidly design gRNAs for gene targets across the genome from a degenerate protospacer adjacent motif (PAM) and a GenBank file. The tool applies Hierarchical Navigable Small World (HNSW) graphs to speed up the comparison of guide RNAs. This allows the user to design gRNAs targeting all genes in a typical bacterial genome in about 1-2 minutes.

**Conclusions:** Guidemaker enables the rapid design of genome-wide gRNA for any CRISPR-Cas enzyme in non-model organisms. While GuideMaker is designed with prokaryotic genomes in mind, it can efficiently process smaller eukaryotic genomes as well. GuideMaker is available as command-line software, a stand-alone web application, and a tool in the CyCverse Discovery Environment. All versions are available under a Creative Commons CC0 1.0 Universal Public Domain Dedication.

## Introduction

CRISPR-Cas technology enables rapid and efficient genome editing in both prokaryotic and eukaryotic cells [1,2]. CRISPR-based systems are set apart from other genome editing tools by the ease with which they can be programmed to target specific sequences. Almost any DNA sequence in the cell can be targeted as long as it possesses a compatible protospacer adjacent motif (PAM). The PAM is a conserved sequence that flanks the DNA target site, known as the protospacer, and must be present for target recognition [3]. The target specifying guide-RNA (gRNA) can be supplied as RNA, or encoded in DNA, depending on the organism under investigation. Although CRISPR-Cas is often used to edit single genes in eukaryotes, it is increasingly used for other purposes in prokaryotic and eukaryotic organisms, including non-model organisms [4].

The *Streptococcus pyogenes* Cas9 (SpCas9) was the first Cas described [5] and it is still the most widely used enzyme in CRISPR in gene editing. Other Cas enzymes described early in the CRISPR revolution, such as the *Staphylococcus aureus* Cas9 and the *Acidaminococcus* Cas12a, are also commonly used [6,7]. Accordingly, the parameters for these enzymes are often included in computational tools to identify CRISPR target sites [8–11]. Cas9 enzymes from other organisms and other Cas-associated proteins that can cleave dsDNA, ssDNA, ssRNA, and insert transposon elements have also been described and have their place in molecular toolkits [12–18]. Each of these enzymes generally possesses its own requirements, such as PAM sequence constraints, PAM orientation, and protospacer length. Many of these CRISPR-Cas systems have been repurposed to enable molecular genetics techniques like gene deletions, gene insertions, transcriptional depletion and activation, and translational repression [12,19–22]. Some of these techniques can be scaled to the genome level with chip-synthesized oligonucleotides and pooled approaches to screening [23]. In pooled screens, high-throughput DNA sequencing is used to identify how the pool has changed over time to elucidate genes that affect cells’ fitness in specific conditions. Given the diversity of the CRISPR systems and their uses, identifying appropriate target sites is not trivial, especially for the number of targets needed for genome-scale experiments.

Here we introduce GuideMaker, a computational tool to identify target sites and design gRNA sequences that is not limited to any specific CRISPR system or organism. Guidemaker is most useful for a few kinds of CRISPR experiments. The first use case is designing pools of gRNAs for genome-wide screening experiments like Perturb-seq and CRISPR pool [23,24]. GuideMaker is optimized for making the all-versus-all comparisons necessary to design a genome-wide screen and return candidate gRNAs for every gene locus. The tool allows the user to filter targets based on their proximity to features of interest, like the start codon for any coding sequence. The second major use case is for researchers working with non-model organisms. Online gRNA design tools often have a limited number of preselected genomes available for analysis because most methods require PAM site positions to be precomputed. GuideMaker rapidly computes all guide positions on demand so the user can provide a set of GenBank files from any organism for analysis. The third use case is for researchers working with Cas enzymes other than the canonical versions of Cas9, Cas12a (Cpf1), or Cas13 with different PAM and target site requirements. GuideMaker allows the user to specify a custom PAM with variable length, including degenerate nucleotides and allows the PAM to be on either the 3’ or 5’ side of the protospacer. These features allow GuideMaker to support any current or future CRISPR-Cas system. Since the determination of which CRISPR-Cas system functions best in any given organism is not predictable, this tool is highly relevant to researchers developing CRISPR tools in new species. In some cases, GuideMaker may not be the best choice. There are mature tools for designing gRNAs in model organisms with common CRISPR-Cas systems and targeting a small number of loci [25,26]. Some of these employ sophisticated statistical models to select the best Cas9 gRNA candidates and may be a better choice for well-studied systems [8]. Because there is limited experimental data on most Cas/organism combinations, GuideMaker relies on design heuristics rather than machine learning-based identification methods.

## Methods

### Main features, input parameters, and workflow

GuideMaker is designed to be easy to use as either a web application or a command-line utility. The key features of GuideMaker are:

1. All the potential guides in a genome can be quickly designed in one run.
2. It can design gRNAs for any small to medium size genome (up to about 500 megabases).
3. It can design gRNAs for any PAM sequence from any Cas system.
4. Search is customizable through user-defined guide parameters (as highlighted in Figure 1). These features are specific to organisms, CRISPR-Cas systems, and experiments. Tuning these parameters can improve the sensitivity and specificity of gRNA.
5. Users can exclude specific restriction sites from guides to preserve those sites for downstream experiments.
6. It creates control gRNAs based on the input genome. In CRISPR experiments it is often desirable to create negative control gRNAs to evaluate off-target binding. GuideMaker provides the user with realistic control gRNAs that are highly divergent from sequences adjacent to PAM sites.
7. Provides an interactive visualization and exploratory tool to evaluate the guides.
8. Provides tabular result files which can be used for the design and ordering of gRNAs.
9. The software can be run as a web application [27], a CyVerse application, or a command-line application [28]. Server code is included for running local instances of the web application as well.

**Figure 1.**
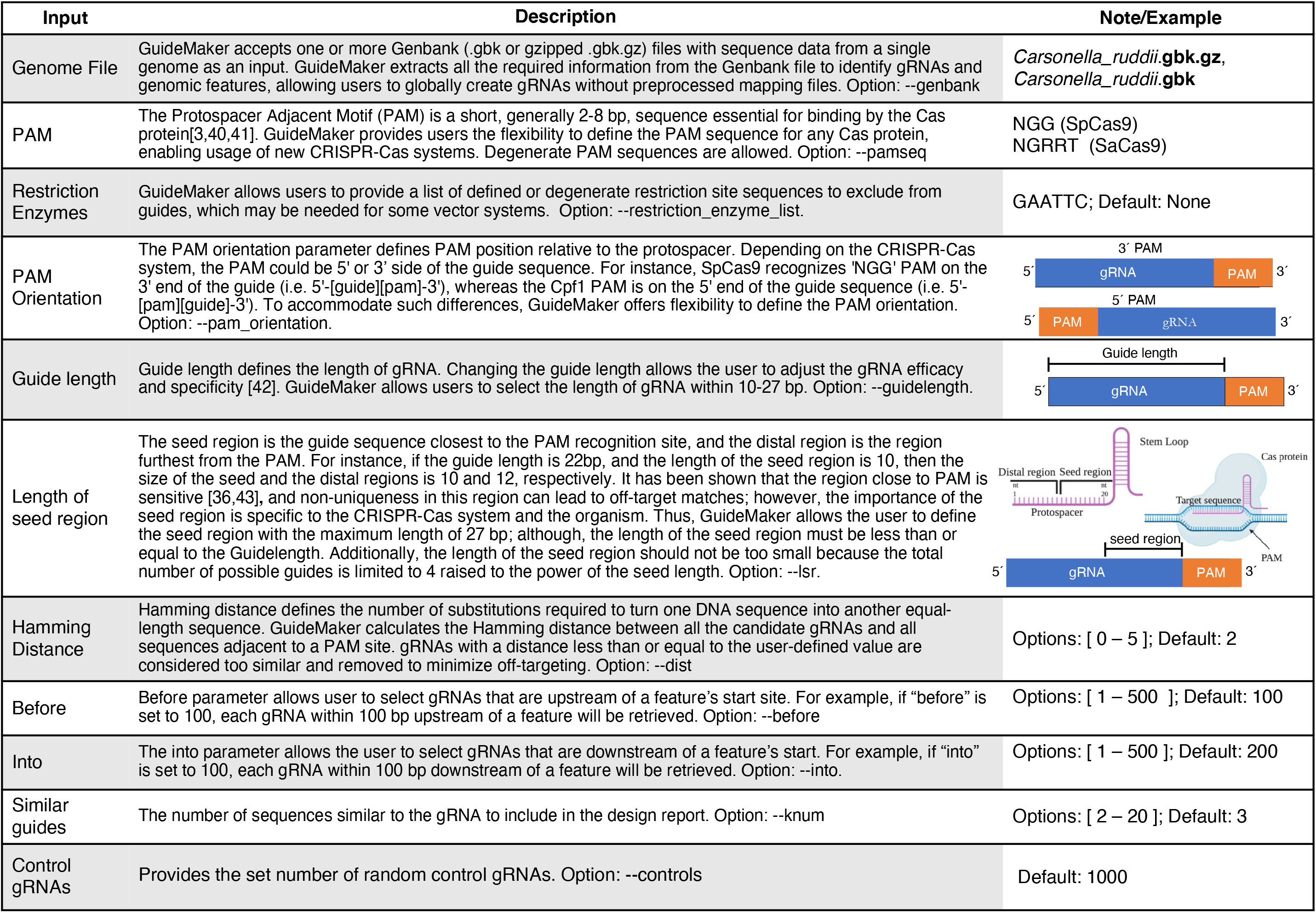
Input parameters for GuideMaker.

A typical workflow of GuideMaker involves three major steps (Figure 2). In the first step, the user uploads the input genome in one or more .gbk or gzipped .gbk files and defines the PAM and gRNA parameters (as highlighted in Figure 1). Guidemaker identifies and filters target sites, then returns summary data to the graphical environment (Figure 2). Users can use the interactive plots to learn more about the identified gRNAs and sort them by genome coordinates or locus tag. In the final step, GuideMaker provides the results as downloadable files under the results section. These files are used for synthesizing guides. The command-line version of GuideMaker has similar input parameters as the web application, with the flexibility to generate plots and configure the underlying hyper-parameters for the Hierarchical Navigable Small World (HNSW) graph, or to run the web application locally. To make the application easier to install we distribute the application as a Bioconda environment [29], Docker container [30], Python package on Github [28], through the Cyverse discovery environment [31] or as an online web application [27]. Detailed information on accessing the software through various methods is available on the project homepage [32].

**Figure 2.**
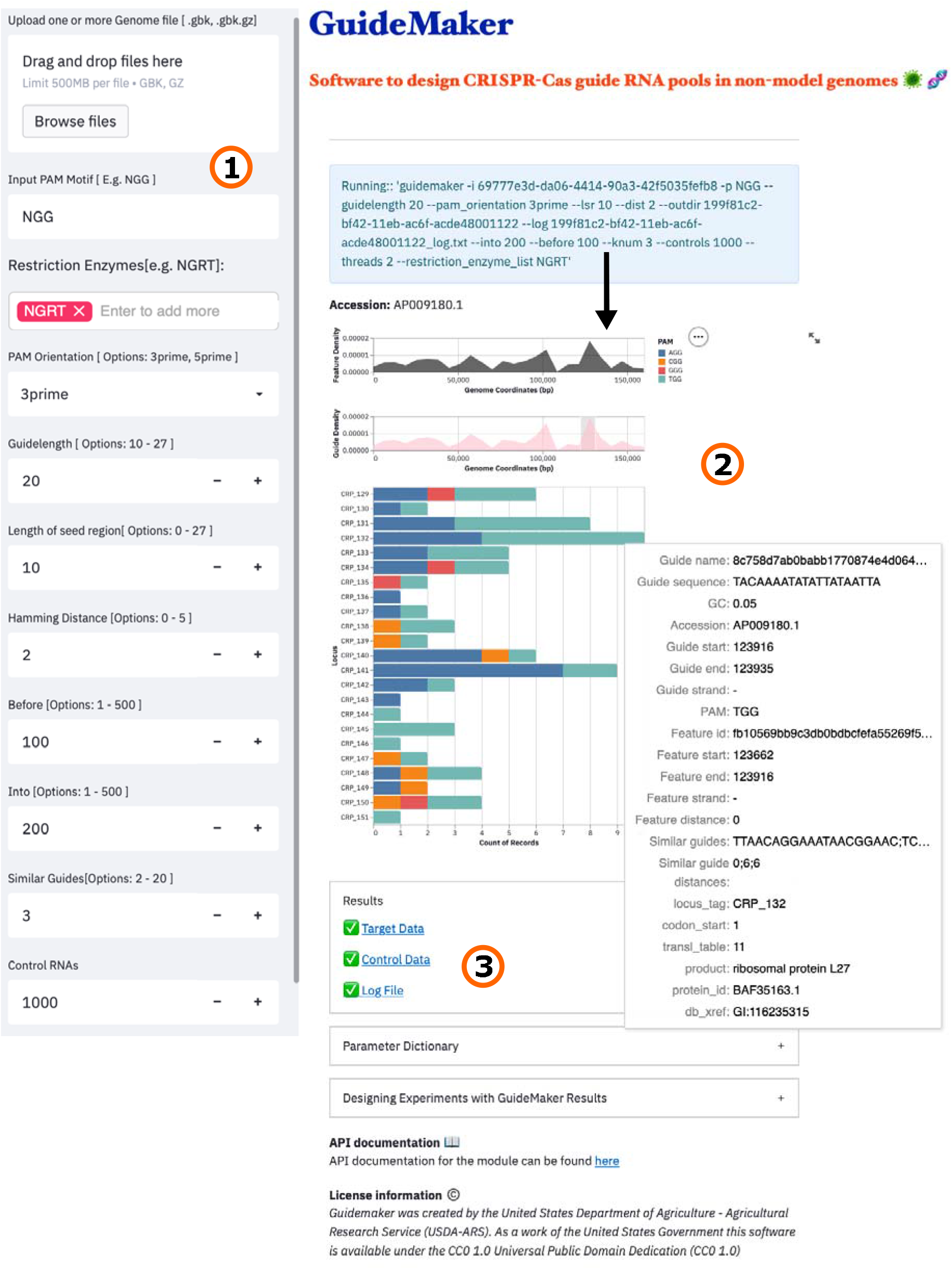
A typical workflow of GuideMaker: 1) A user uploads the input genome (single or multiple) as Genbank file, then defines the PAM sequence along with all the associated parameters and submits them to run the program. 2) GuideMaker processes the input files and generates the interactive plots. Users can use these interactive plots to explore the results and sort them by locus tag and genome coordinates. 3) GuideMaker provides all the results and log files as downloads under the “Results” section.

### Search method

GuideMaker initially scans the genome, recording all candidate guide sequences adjacent to the specified PAM sequence on both DNA strands (Figure 3). Candidate guides are then optionally checked for the restriction sites. Next, the candidates guides are searched for a unique “seed region” closest to the PAM site and candidate gRNAs that are not unique in their “seed region” are removed. Then, approximate nearest neighbor search is used to remove candidate guides too similar to PAM adjacent sequences in the genome, based on Hamming distance (the number of substitutions required to turn one DNA sequence into another equal-length sequence). The approximate nearest neighbor search is performed using the Hierarchical Navigable Small World (HNSW) graph method in the Non-Metric Space Library (NMSLIB) [33,34]. An index of all the initial candidate guides is created using the bitwise Hamming distance metric. Each guide with a unique “seed region” is compared to all candidate guides and any guides with Hamming distances below the user-set threshold are removed. This differs from the standard procedure of indexing the genome and mapping each candidate guide against the whole genome then parsing each result. HNSW has a search complexity of 𝒪 (log *N*) and index complexity of 𝒪 (*N* · log *N*)[33]. Finally, user-defined criteria are applied specifying the proximity and orientation of guides relative to genomic features like genes. A list of guides is then returned to the user with relevant information about the guide and its target genomic features.

**Figure 3.**
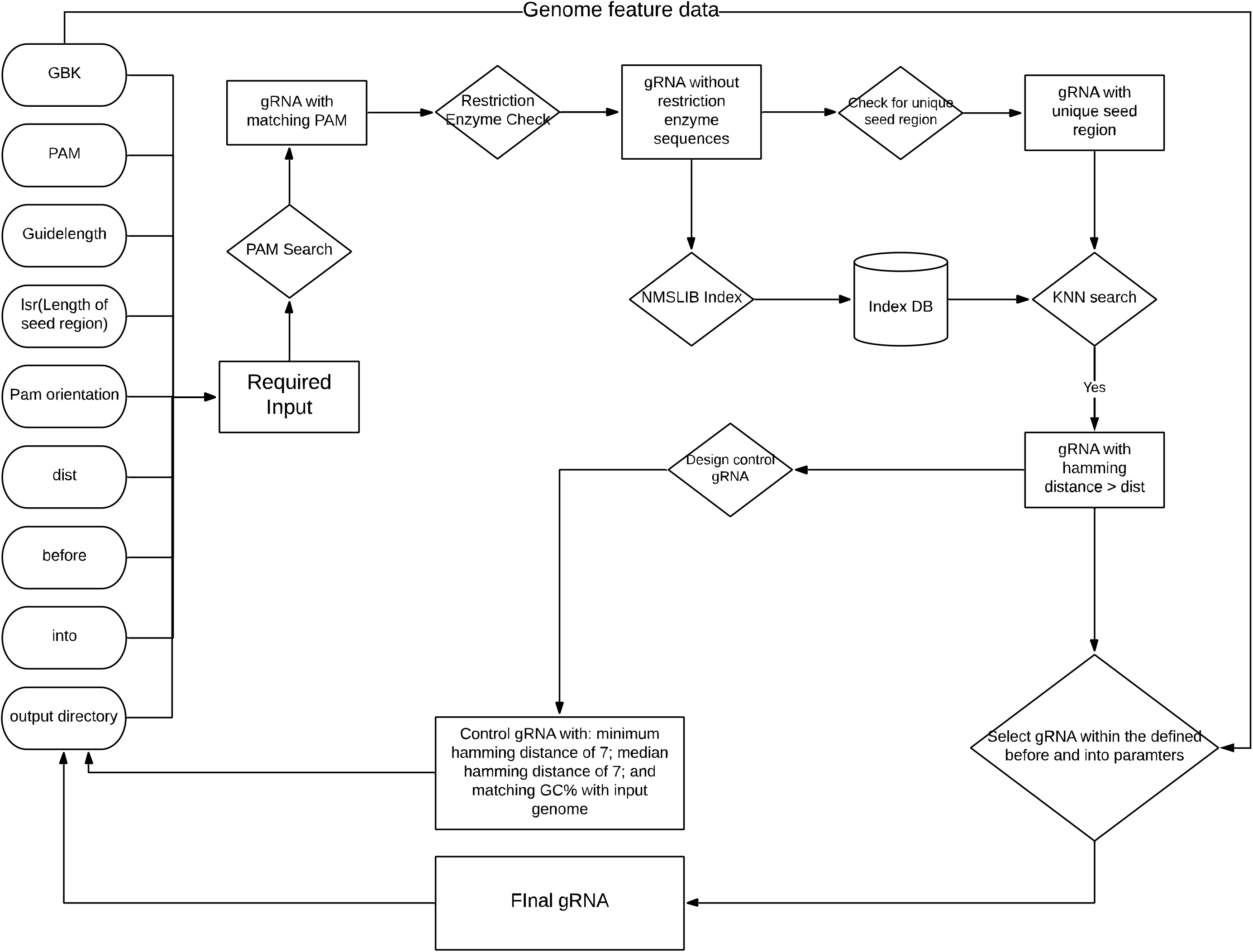
Entity Relationship Diagram showing the operation of the GuideMaker core program.

The core of GuideMaker’s search method is the HNSW method in NMSLIB [34]. The method builds a multilayer graph index of the input data and has several parameters that can be optimized for index building and search to trade-off speed and accuracy. Graph construction is the most time-consuming step in our tests, and thus grid optimization was run to minimize run time while keeping recall above 99% relative to the ground truth exact nearest-neighbor search. The grid-optimization parameters: [M, efc, ef, and post] used in the HNSW graph for approximate nearest neighbor search have been optimized for bacterial genomes. A script for re-optimization (flag --config) of these hyper-parameters is included in the command-line version of the software.

### Computational performance

Genomes of different sizes, GC content, and chromosome numbers were used to test the speed and scalability of GuideMaker (Supplementary Table 1). For benchmarking the performance, the same parameters were used unless a specific parameter was being tested: a PAM motif of ‘NGG’, 3’ pam orientation, target length of 20, lsr (length of seed region) of 11, before and after parameters of 500, knum of 10, controls of 10, dist of 3 and threads of 32. We profiled the performance of GuideMaker with different threads [1, 2, 4, 8, 16, and 32] in processors with and without the AVX2 processor instruction set. All tests were run on a single compute node with 2 x 24 core Intel(R) Xeon(R) Platinum 8260 CPU @ 2.40 GHz with Cascade Lake microarchitecture. Three bacterial genomes, a fungal genome, and a plant genome were used in performance benchmarking: *Escherichia coli* (K12), *Pseudomonas aeruginosa* (PAO1), *Burkholderia thailandensis* (E264), *Arabidopsis thaliana*, and *Aspergillus fumigatus*. For the gene or locus-specific comparisons, only the guides within the locus coordinates (i.e. zero feature distance) were considered.

### Comparison to existing design method

We compared the results of GuideMaker with the results of the online version of CHOPCHOP[35]. GuideMaker and CHOPCHOP parameters were set to approximate the same search. The length of the target sequence was set to 20 and zero mismatches were allowed in the seed region (11bp) of the target. The *Escherichia coli* (str. K-12/MG1655) genome was used with the online version of CHOPCHOP since it has a limited number of genomes. Targets were searched in 40 Kbp increments to account for CHOPCHOP’s size limitations. Target sequences were searched across multiple 40 Kbp segments of *E.coli* genome (NC_000913.3:2001-42000, NC_000913.3:80001-120000, NC_000913.3:160001-200000, NC_000913.3:240001-280000, and NC_000913.3:320001-360000). We also searched for target sequences and genes/locus_tags within 40Kbp of (NC_000913.3:2001-42000) to compare identifications at the locus level. Ratios between tools were calculated by dividing the number of gRNA identify with GuideMaker by the number of CHOPCHOP identified gRNA to represent the proportion of guides identified by both GuideMaker and CHOPCHOP.

## Results

The time for Guidemaker to complete a typical run identifying all SpCas9 gRNAs (PAM ‘NGG’) in a bacterial genome using 8 compute cores was 75 seconds for *E. coli* and 130 seconds for *P. aeruginosa* (Figure 4). For SaCas9 and StCas9, which have a longer PAM sequence (‘NGRRT’ and ‘NNAGAAW’ respectively, with 3’ PAM orientation) and thereby fewer potential targets, the same genomes ran in 19 or 5 seconds (Supplementary Figures 1). The fungus *Aspergillus fumigatus* (28MB) and plant *Arabidopsis thaliana* (114 MB) have larger genomes but are still processed quickly. *A. fumigatus* processed between 23 – 304 seconds, while *A. thaliana* processed in 250-921 seconds depending on the number of cores, AVX2 instructions, and PAM sequence (Supplementary Figures 2). GuideMaker can take advantage of Advanced Vector Extensions (AVX2) on newer x86 processors, which improves the search speed because HNSW search is accelerated with AVX2 (Supplementary Figure 3). The acceleration was larger when fewer processors were available (Supplementary Figure 3). With more processors, the run time was similar regardless of AVX2 use. The HNSW algorithms are parallelized, and indexing-and-search takes most of the compute time in GuideMaker so the software scales well when additional cores are added up to 8 cores (Supplementary Figure 3). In practice it scaled up sub-linearly with genome size, globally estimating Cas9 guides for *E. coli* MG1655 (4.6MB) in 75 seconds and *A. thaliana* (114.1MB) in 921 seconds, both on 8 cores (Memory usage: 1.9GB for E. coli and 15.4 GB for *A. thaliana*, Supplementary Figure 4).

**Figure 4.**
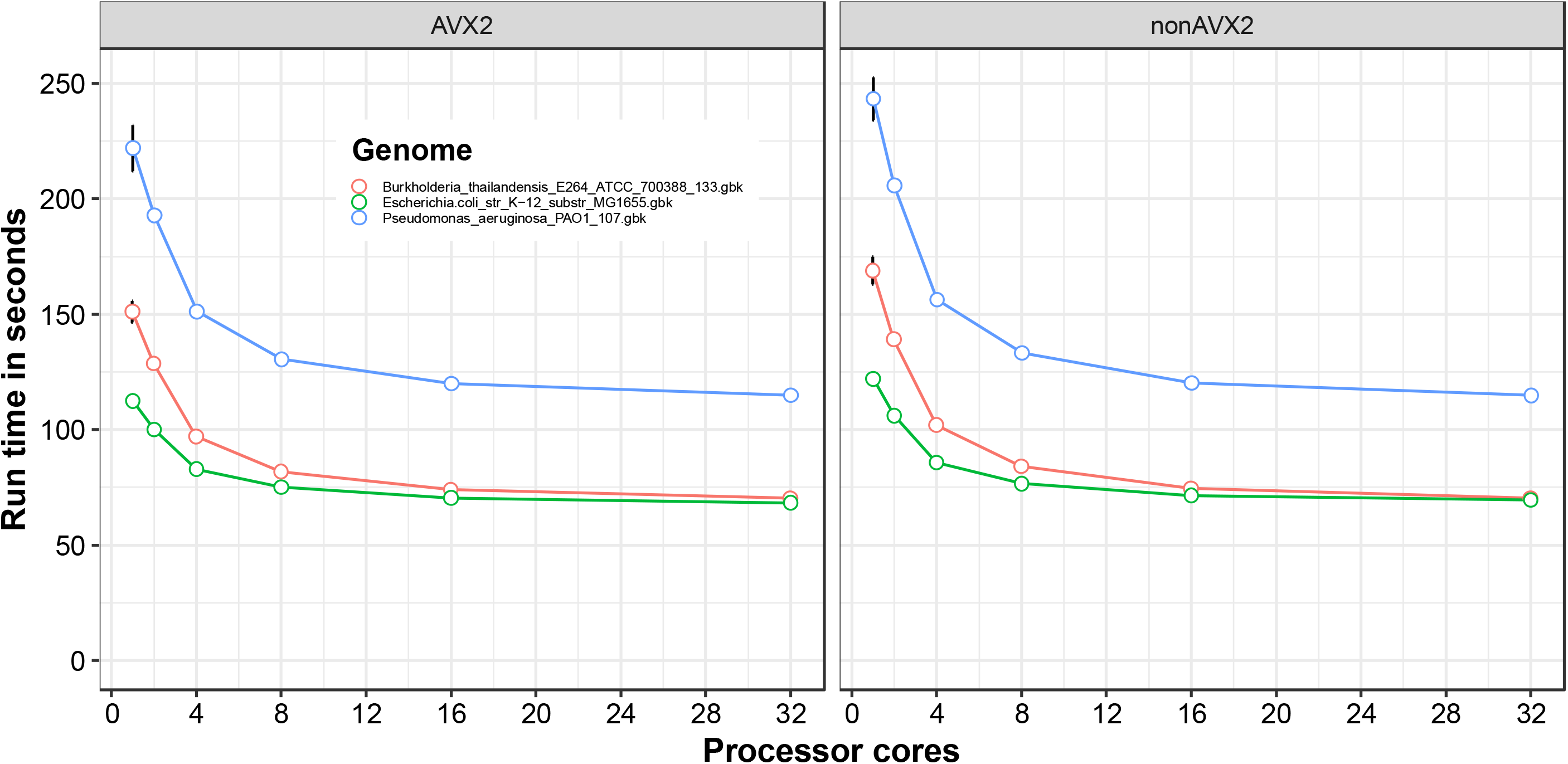
Performance of GuideMaker for SpCas9. Evaluating the performance of GuideMaker across three bacterial genomes using the **“NGG”** PAM motif with a target length of 20, unique zone of 11, 3prime PAM orientation, before and into parameters of 500, knum of 10, controls of 10, and dist of 3. The mean of 10 runs was used for the evaluation, where dot and bar represent the mean and standard error, respectively.

The results of Guidemaker were compared with the popular guide design software CHOPCHOP version 3 [35]. When GuideMaker’s filtering settings are set to match CHOPCHOP, the results are very similar and 99.9% of the targets identified by GuideMaker fall within 2bp of target coordinates returned by CHOPCHOP. When GuideMaker’s unique seed region criterion was not applied at the loci level, the average number of guides identified by the two approaches was similar per locus (Mean GuideMaker = 116.8, Mean CHOPCHOP = 113.6, p-value = 0.86, Supplementary Table 2). Although the number of guides identified per gene locus differed, none of the genes were missed by either tool. GuideMaker’s default requirement of a seed region is more stringent than CHOPCHOP, and with it enabled, GuideMaker returns (count=1787) 38.4% (for 2Kbp-42Kbp regions) of the targets compared to CHOPCHOP (count=4651) *E. coli* K12. At the sequence level, 96.7% of the identified gRNA (1729/1787) from both of the tools had identical sequences. The more stringent filtering could potentially reduce off-targeting but that would need to be experimentally validated in a range of organisms. The ratio of gRNA found by both the tools across the multiple 40Kbp regions was 39.2% (sd= 1.9%, Supplementary Table 3) when using Guidemaker’s more stringent default settings. This ratio was calculated by dividing the number of gRNA from GuideMaker by the number from CHOPCHOP for each 40Kb region.

## Discussion

Designing gRNAs is a two-step process where GuideMaker first identifies potential guides adjacent to PAM sequences and then filters the potential guides based on multiple criteria. The most important criterion is that each guide has a minimum edit distance from any other sequence adjacent to a PAM site in the genome; this decreases the likelihood of off-target binding. The second way GuideMaker reduces off-target binding is by requiring that a set number of bases near the PAM site are unique from any other candidate guide. The 8 bases nearest the PAM are the most important for target specificity, and any mismatch is sufficient to prevent binding [36,37]. The length of the unique region should be set with consideration for the size of the genome since requiring short unique regions will limit the number of total guides that can be found. For example, requiring that every gRNA be unique in the first 3 bp would only allow for 4^3^ = 64 possible guides to be designed. For normal *--lsr* values of 9-12 this is only limiting for human-sized genomes and can be disabled by setting *--lsr* to 0. All guides designed by GuideMaker are perfect matches to a single site in the genome. Specificity is obtained by requiring all similar PAM-adjacent sequences to be unique in the critical “seed region” and have a total number of mismatches that exceed the user-defined threshold. This double criterion is expected to increase specificity.

The primary goal of the current version of our software is to support the design of gRNAs in non-standard Cas enzymes for non-model organisms at the genome-scale. It is known that gRNA’s do not perform equally, thus empirical experiments will be needed to fully validate the functionality and efficacy of gRNA predictions. Given the similarity in targets identified by GuideMaker and CHOPCHOP, we anticipate performance will be similar to the current state of the art but applicable to more design use cases. When a unique seed region and Hamming distance-based filters were applied, GuideMaker created guides more conservatively, generating only about 40% of the guides created by CHOPCHOP. While CHOPCHOP has an option to specify the maximum number of mismatches in the first 9 bp or the whole guide, it does not allow the application of both criteria. While there are small differences in the number and position of guides generated by GuideMaker, with GuideMaker being more conservative by default, both programs create enough guides to target nearly all gene loci in the genome of *E. coli*. If experimentally validated data become available from genome-wide screens with different Cas enzymes, the future versions of GuideMaker could potentally incorporate scoring matricies to help rank candidate guides.

Guidemaker is a fast and flexible tool for designing guide RNA across the entire genome in non-model organisms or with non-canonical Cas enzymes. It takes advantage of fast HNSW search to quickly index and search new genomes. Several parameters can be tuned to ensure compatibility with the specific application of the user. For example, GuideMaker checks the designed gRNA for a given restriction enzyme site to prevent incompatibility with the cloning strategy. Second, the maximum distance from a target sequence from the start of an annotated feature can be chosen to disrupt promoters or the beginning of the coding sequence, since these sites are preferred for CRISPRi experiments. GuideMaker also creates off-target gRNAs for use as negative controls in high-throughput experiments. Lastly, the program plots the results for visual exploration of the targets and exports the data as .csv files. The software is available as a command-line application, a web application, and is integrated into the CyVerse Discovery Environment to provide users with a range of usage options.

## Availability and Requirements

Project name: GuideMaker

Project home page: https://guidemaker.org

Operating system(s): Linux or MacOS Programming language: Python >=3.6

Other requirements: ‘pybedtools==0.8.2’, ‘nmslib>=2.0.6’,’altair’, ‘streamlit>0.80.0

License: CC0 1.0 Public Domain Dedication

## Competing Interests

Authors declare no competing interests

## Data Availability

The source code and command-line executables for GuideMaker are available at the Zenodo [38] and can be installed directly from Github [28], Bioconda [29], or as a Docker container [30]. Data and code to reproduce the analysis in the paper are available at Zenodo [39]. As a work of the United States Department of Agriculture, Guidemaker is released to the public domain under a Creative Commons (CC0) public domain attribution. The program is also available as a web application through the Cyverse discovery environment [31], and as a stand-alone web application [27].

## Additional Files

**Supplementary Figure 1**. Performance of GuideMaker for SaCas9 and StCas9.

**Supplementary Figure 2**. Performance of GuideMaker for SpCas9, SaCas9, and StCas9.

**Supplementary Figure 3**. Performance of GuideMaker with AVX2 settings.

**Supplementary Figure 4**. Memory usage of GuideMaker for SpCas9, SaCas9, and StCas9.

**Supplementary Table 1:** Organism features

**Supplementary Table 2:** Comparison of the average number of gRNA identified by GuideMaker and CHOPCHOP.

**Supplementary Table 3:** Comparison of consensus ratio between GuideMaker and CHOPCHOP.

## List of abbreviations

CAS: CRISPR-associated protein
CRISPR: Clustered Regularly Interspaced Short Palindromic Repeats
gRNA: Guide RNA
HMSW: Hierarchical Navigable Small World
NMSLIB: Non-Metric Space Library
PAM: Protospacer Adjacent Motif

## Funding

The research was supported by the United States Department of Agriculture (USDA), Agricultural Research Service (ARS) project number 6066-21310-005-D, and ARS cooperative agreement 6066-21310-005-28-S to the University of Florida. This research used resources provided by the SCINet scientific computing initiative of the USDA-ARS, ARS project number 0500-00093-001-00-D.

## Author Contributions

R.P., L.T.R., C.R.R., and A.R.R. conceived and designed the study. R.P. and A.R.R developed and optimized the software and performed the experiments. R.P., L.T.R., C.R.R., and A.R.R, tested the software, wrote, and revised the manuscripts. All authors read and approved the final manuscript.

## Notes

https://guidemaker.org

### Competing Interest Statement

The authors have declared no competing interest.

