## Supplemental Data for "GuideMaker: Software to design CRISPR-Cas guide RNA pools in non-model genomes"

**Supplementary Materials**

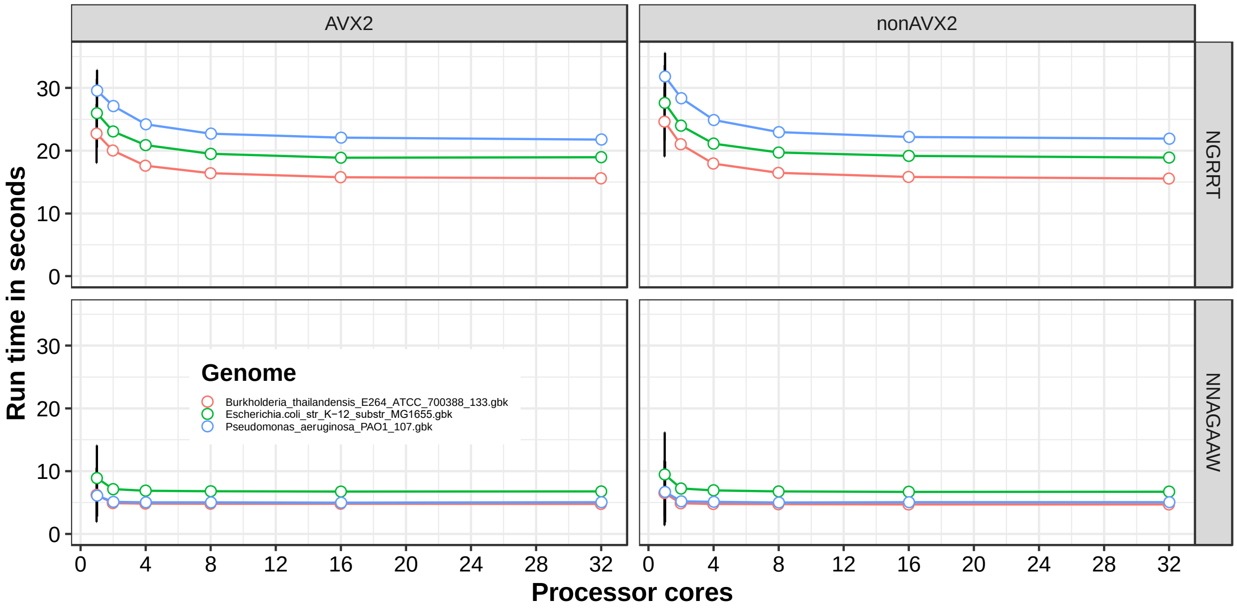


**Supplementary Figure 1. Performance of GuideMaker for the Cas enzymes SaCas9 and StCas9.** Evaluating the performance of GuideMaker across three bacterial genomes using the **"NGRRT"** and the **"NNAGAAW"** PAM motifs with target length of 20, unique zone of 11, 3prime pam orientation, before and into parameters of 500, knum of 10, controls of 10, and dist of 3. The mean of 10 runs was used for the evaluation, where dot and bar represent the mean and standard error respectively.

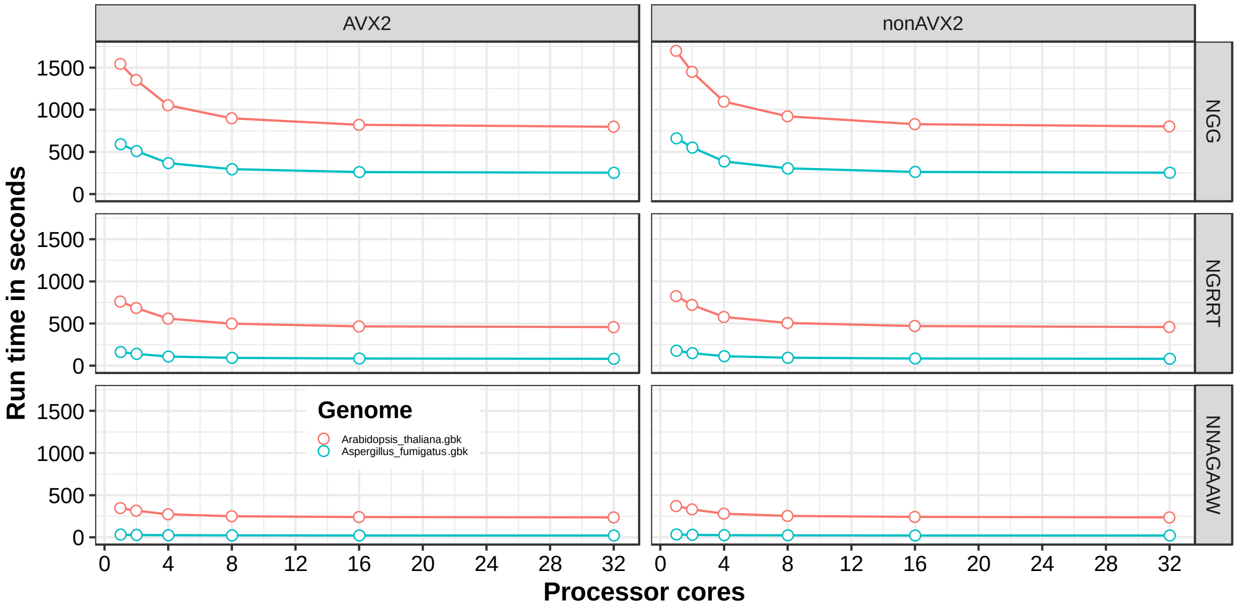


**Supplementary Figure 2. Performance of GuideMaker for SpCas9, SaCas9 and StCas9**.

Evaluating the performance of GuideMaker in larger genomes: *Arabidopsis thaliana* (~ 114.1MB) and *Aspergillus fumigatus* (28 MB) using the **"NGG"**, the **"NGRRT"** and the **"NNAGAAW"** PAM motifs with target length of 20, unique zone of 11, 3prime pam orientation, before and into parameters of 500, knum of 10, controls of 10, and dist of 3. The mean of 10 runs was used for the evaluation, where dot and bar represent the mean and standard error respectively.

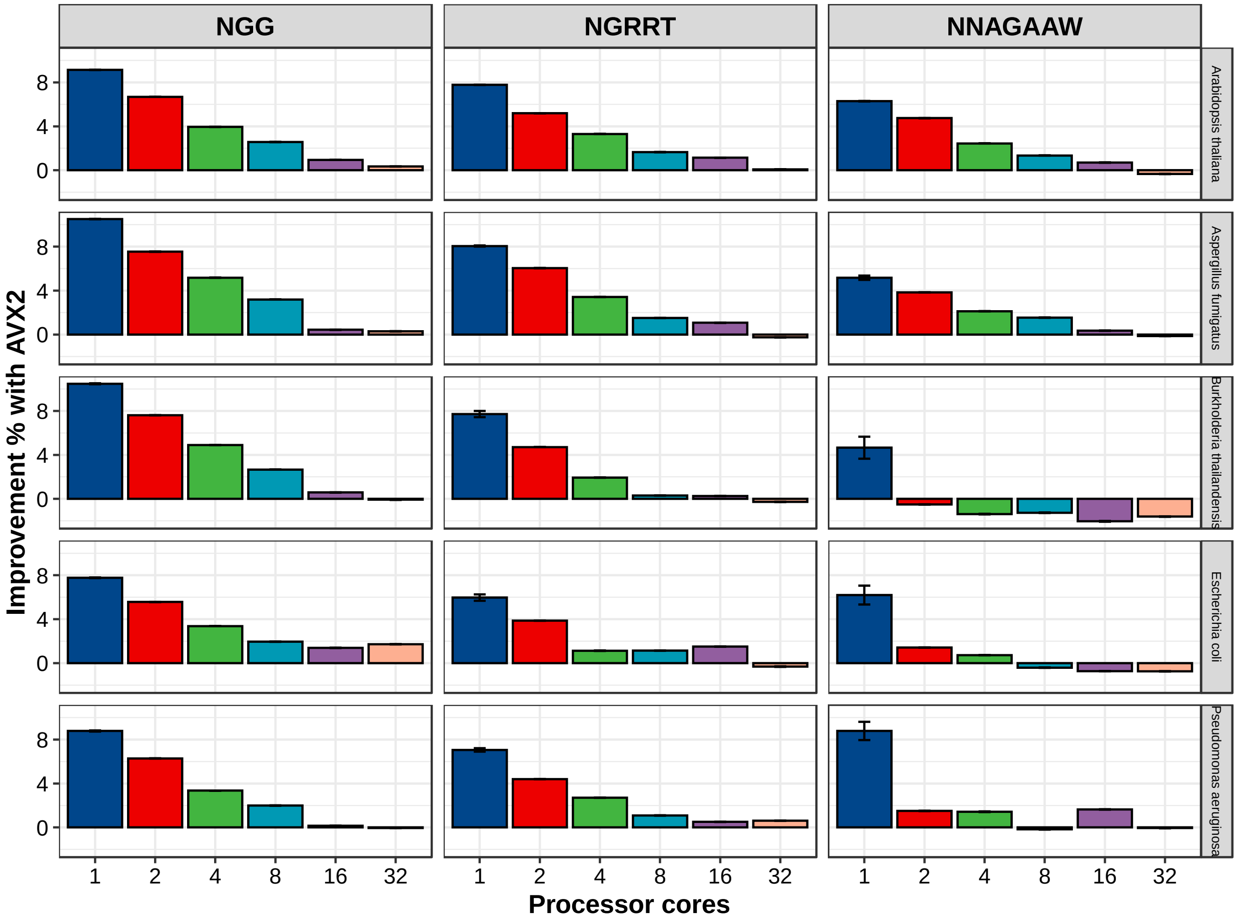


**Supplementary Figure 3. Performance of GuideMaker with AVX2 settings.** The speedup of Guidemaker wit haVX2 was compared with different numbers of logical cores (1, 2, 4, 8, 16, and 32) respectively. GuideMaker run times for 5 different genomes (*Escherichia.coli_str_K-12_substr_MG1655, Pseudomonas_aeruginosa_PAO1_107, Burkholderia_thailandensis_E264_ATCC_700388_133, Aspergillus fumigatus, and Arabidopsis thaliana*) with three different PAM (**"NGG"**, **"NGRRT"** and **"NNAGAAW"**) across 10 runs were used to calculate Improvement % with AVX2 instruction. Height of the bar represents Improvement % with AVX2 instruction and the error bar represents the standard deviation.

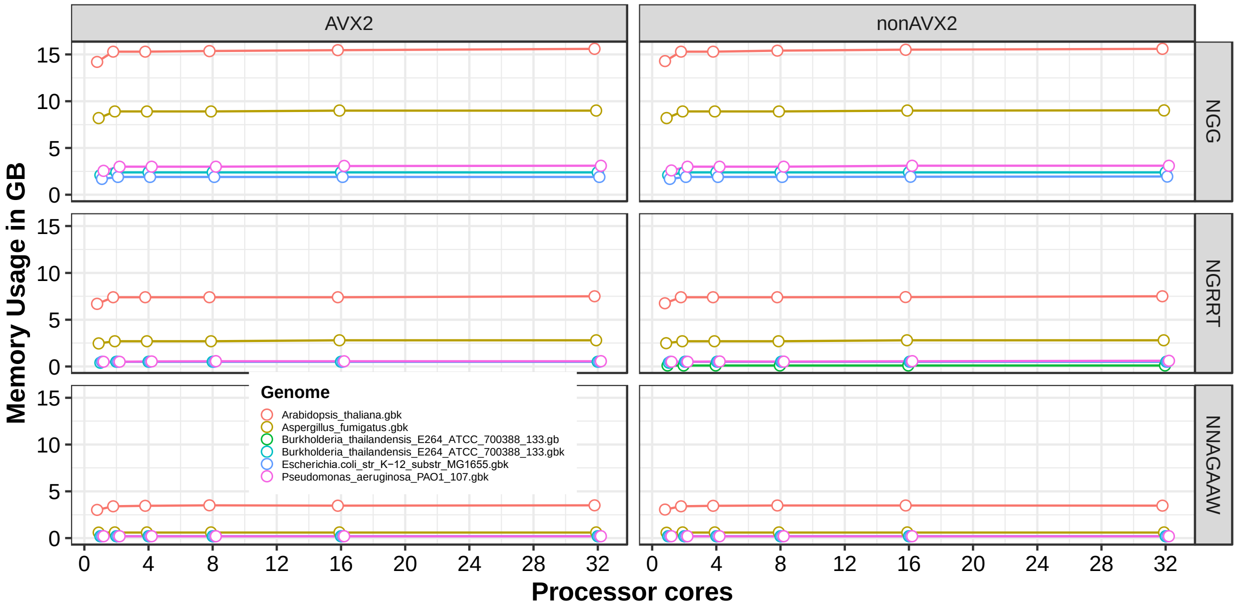


**Supplementary Figure 4. Memory usage of GuideMaker for SpCas9, SaCas9, and StCas9**. Evaluating the memory usage of GuideMaker when the different number of logical cores with or without AVX2 settings are available. All the genomes were evaluated using the same parameters for GuideMaker (target length of 20, unique zone of 11, 3prime pam orientation, before and into parameters of 500, knum of 10, controls of 10, and dist of 3) with three PAM motifs ("NGG", "NGRRT", and "NNAGAAW"). The mean of 10 runs was used for the evaluation, where dot and bar represent the mean and standard error respectively.

**Supplementary Table 1:** Organisms features

| Organisms | Genome Size (MB) | GC % | Locus |
|---|---|---|---|
| Escherichia.coli_str_K-12_substr_MG1655 | 4.4 | 50.79 | 4357 |
| Pseudomonas_aeruginosa_PAO1_107 | 6.0 | 66.56 | 5584 |
| Burkholderia_thailandensis_E264_ATCC_700388_133 | 6.4 | 67.63 | 5633 |
| Aspergillus fumigatus | 28.0 | 48.2 | 9630 |
| Arabidopsis thaliana | 114.1 | 36.0 | 48262 |

**Supplementary Table 2:** Comparison of an average number of gRNA predicted by GuideMaker and CHOPCHOP.

| | | | | Welch Two Sample t-test | | |
|---|---|---|---|---|---|---|
| | Mean | N | sd | t | df | p-value |
| GuideMaker | 116.89 | 37 | 83.66 | 0.1675 | 71.949 | 0.8674 |
| ChopChop | 113.67 | 37 | 81.46 | | | |

Comparison of an average number of gRNA predicted by GuideMaker (without applying unique seed-region based and hamming distance-based filters) and CHOPCHOP for all the genes located within 2Kbp-42Kbp region of *E. coli* MG1655.

**Supplementary Table 3:** Comparison of consensus ratio between GuideMaker and CHOPCHOP.

| Genome Coordinate | GuideMaker | CHOPCHOP | Ratio | Average Ratio |
|---|---|---|---|---|
| NC_000913.3:2001-42000 | 1787 | 4651 | 0.38 | 0.392 |
| NC_000913.3:80001-120000 | 1856 | 4619 | 0.40 | |
| NC_000913.3:160001-200000 | 1718 | 4663 | 0.37 | |
| NC_000913.3:240001-280000 | 2003 | 4819 | 0.42 | |
| NC_000913.3:320001-360000 | 1856 | 4730 | 0.39 | |

Ratio = Number of gRNA predicted by GuideMaker / Number of gRNA predicted by CHOPCHOP

Comparison of consensus ratio between GuideMaker and CHOPCHOP using multiple genome segment and genes in *Escherichia coli* (str. K-12/MG1655). The ratio between GuideMaker and CHOPCHOP was calculated as the proportion of guides produced by both the tools.
